# BioID based proteomic analysis of the Bid interactome identifies novel proteins involved in cell cycle dependent apoptotic priming

**DOI:** 10.1101/529685

**Authors:** Robert Pedley, Louise E. King, Venkatesh Mallikarjun, Pengbo Wang, Joe Swift, Keith Brennan, Andrew P. Gilmore

## Abstract

Apoptotic priming controls the commitment of cells to apoptosis by determining how close they lie to mitochondrial permeabilisation. Variations in priming are important for how both healthy and cancer cells respond to chemotherapeutic agents, but how this is dynamically coordinated by Bcl-2 proteins remains unclear. The Bcl-2 family protein Bid is phosphorylated when cells enter mitosis, increasing apoptotic priming and sensitivity to anti-mitotic drugs. Here we report an unbiased proximity biotinylation (BioID) screen to identify regulators of apoptotic priming in mitosis, using Bid as bait. The screen primarily identified proteins outside of the canonical Bid interactome. Specifically, we found that voltage-dependent anion-selective channel protein 2 (VDAC2) was required for Bid phosphorylation dependent changes in apoptotic priming during mitosis. Thus, we identify an additional layer of regulation upstream of known Bid interactions that control dynamic changes in apoptotic priming. These results highlight the importance of understanding the wider Bcl-2 family interactome and its role in regulating the temporal control of apoptotic priming.

## Introduction

Most cells respond to an apoptotic stimulus *via* mitochondrial outer membrane permeabilisation (MOMP) (1). MOMP releases pro-apoptotic factors like cytochrome *c* into the cytosol (2). However, the proportion of cells that undergo MOMP varies both within a cell population as well as between different cell types and tissues (3–5). Apoptotic priming refers to how close a cell lies to MOMP (6, 7). Cells that are “primed” readily undergo MOMP following a pro-apoptotic signal, whereas “unprimed” cells can tolerate a greater level of damage. Determining how primed cancer cells are can predict patient response to chemotherapy (8, 9). Apoptotic priming is dynamic, adjusting as cells response to internal and external conditions, such as survival signals and cell cycle progression (7, 10). How these changes in priming are dynamically coordinated is poorly understood.

MOMP is controlled by the Bcl-2 family of proteins, comprising: pro-apoptotic Bax and Bak, which form the pores that drive MOMP; anti-apoptotic proteins, such as Bcl-XL, Bcl-2, and Mcl-1, that suppress MOMP by sequestering pro-apoptotic proteins; and the BH3-only proteins, which regulate the activity of the other two groups (11, 12). BH3-only proteins may be activators or sensitisers. Activators, like Bim and Bid, may bind Bax and Bak directly to initiate MOMP. These activators can be sequestered by anti-apoptotic Bcl-2 proteins, but are released by sensitisers, such as Bad (13). Thus, variation in the expression and interaction landscape of the different Bcl-2 family subgroups can adjust apoptotic priming, setting the threshold for MOMP. Subtle changes in this balance can have profound consequences (14).

Cells become primed for MOMP in mitosis, with those that fail to divide eliminated (10). Once mitotic cells satisfy the Spindle Assembly Checkpoint (SAC), they enter anaphase and priming is reduced. Drugs like Taxol exploit this checkpoint by preventing transition to anaphase. However, cancer cells can exit mitosis without dividing, termed slippage (15). Slippage occurs if a cell resists apoptosis long enough such that, despite the absence of correct spindle attachment, the gradual degradation of cyclin B1 results in mitotic exit. Increased priming in mitosis has been linked to post-translational modification of several Bcl-2 proteins (16–21), changes in their expression (22, 23) and their degradation (17, 24). Together, these events set the mitotic Bcl-2 landscape and determine how long a cell can resist apoptosis whilst attempting to satisfy the SAC. Thus, mitosis provides an opportunity for understanding how cells dynamically adjust apoptotic priming within precise temporal boundaries.

We have previously demonstrated that the full-length version of the BH3-only protein Bid is phosphorylated on serine 66 during mitosis (21). This phosphorylation increased apoptotic priming specifically in mitosis, increasing cell sensitivity to apoptosis when treated with Taxol. Importantly, this phosphorylation dependent function of full-length Bid did not involve the canonical regulation *via* caspase 8 mediated proteolysis. To understand how cells coordinate temporal changes in apoptotic priming, we have now performed an unbiased proteomic screen based upon proximity-dependent biotin identification (BioID) (25). Using full-length (FL) Bid as bait, we show that the predominant proteins identified in live cells are outside of the canonical Bid interactome and the immediate Bcl-2 protein family. In particular, we identify that the outer mitochondrial porin, VDAC2, is required for cells to temporally increase apoptotic priming in response to mitotic Bid phosphorylation.

## Results

### Using BioID to interrogate the Bid interactome in live cells

In order to interrogate temporal regulation of priming, we decided to utilize post-translational modifications (PTMs) known to alter Bcl-2 protein function in mitosis. Mitosis is particularly useful for a proteomic screen as cells can readily be enriched in M phase. We previously demonstrated that Bid becomes phosphorylated in mitosis, on serine 66 in mouse and 67 in human (mBid and hBid respectively) (21). Bid phosphorylation increased priming in colon carcinoma cells, and thus their sensitivity to apoptosis during Taxol-induced mitotic arrest. We decided to exploit this temporal regulation of Bid phosphorylation to identify mitosis specific regulators.

To identify potential regulators of Bid function in mitosis, we employed a biotin labelling identification (BioID) strategy. BioID is a powerful approach for interrogating protein complexes in live cells (25–28). A bait protein is expressed as a fusion with the *E.coli* biotin ligase BirA containing a R118G substitution along with a myc epitope (from here on referred to as BirA*). On the addition of excess biotin to the culture media, BirA* generates reactive biotinyl-5’-AMP that can covalently bond to primary amines. The half-life of reactive biotinyl-5’-AMP limits the effective labelling radius to ~20 nm (25). Consequently, if BirA* is expressed fused to a bait protein, there will be enrichment for biotinylated proteins proximal to that bait, which can be isolated by streptavidin affinity purification (Fig.1A). As labelling occurs *in situ*, BioID is particularly useful for interrogating complexes that might be sensitive to detergent extraction, a known issue within the Bcl-2 family (29, 30). We therefore generated mBid-BirA* baits to identify interacting partners in live cells.

**Figure 1.**
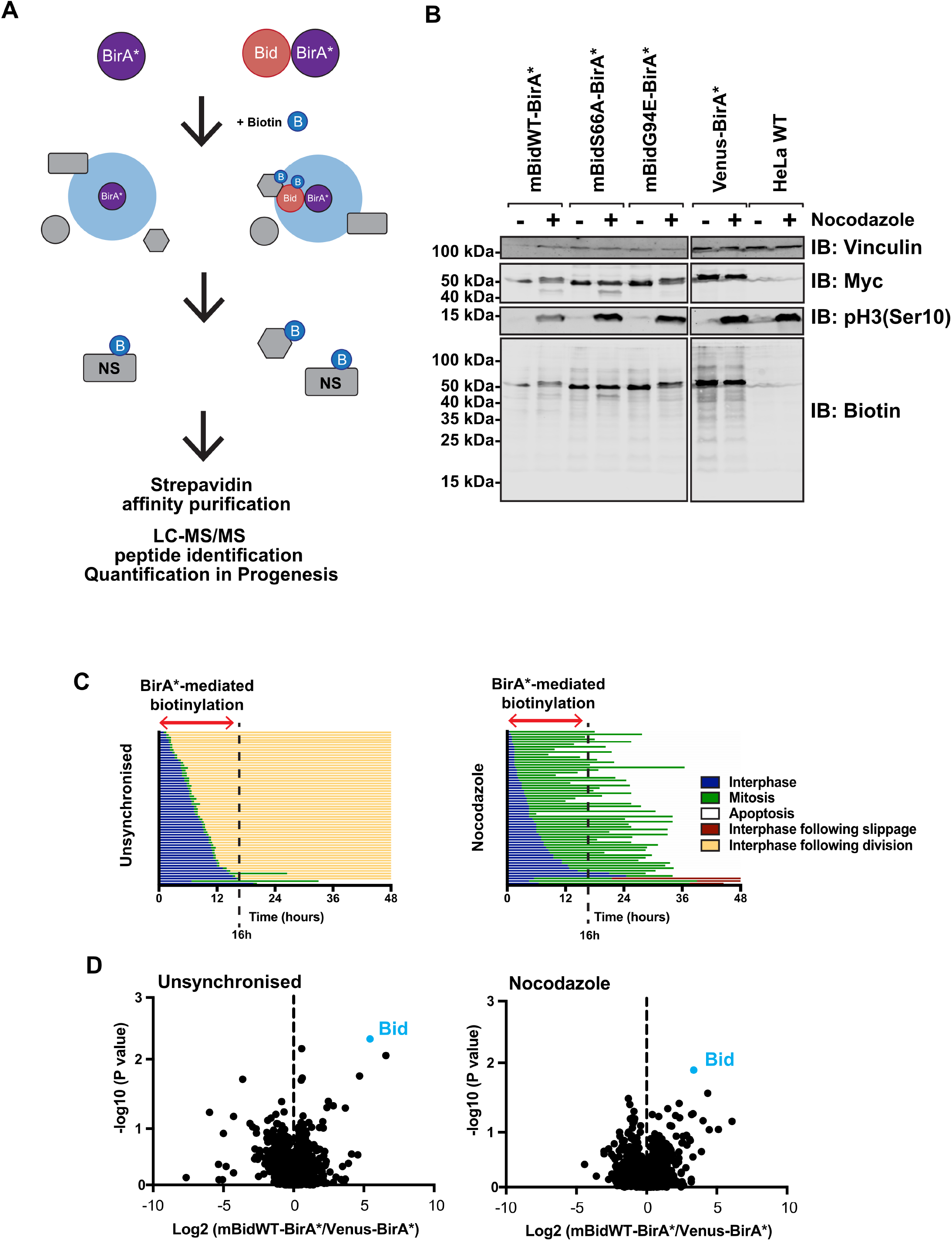
BioID workflow for identification of Bid vicinal proteins in mitotic cells. A. Schematic diagram of BioID labelling strategy. Selective biotinylation of proximal proteins is followed by stringent cell lysis for streptavidin affinity purification and identification by LC-MS/MS. B. HeLa cells, either wildtype or stably expressing mBidWT-BirA*, mBidS66A-BirA*, mBidG94E-BirA* or Venus-BirA*, were grown with 50 μM biotin for 16 hours in the presence (+) or absence (−) of nocodazole. Whole cell lysates were prepared and examined by immunoblotting for the indicted anti-bodies and streptavidin. C. Single cell fate profiles of HeLa cells in the presence or absence of nocodazole, imaged over 48 hours. Each individual line represents a single cell. Data represents 60 cells tracked over two independent experiments. A biotin labelling window of 16 hours significantly enriched for mitotic cells compared to untreated controls, without significant enrichment for apoptotic cells. D. Volcano plot of mean fold change of biotinylated protein abundance for mBidWT - BirA* vs. venus-BirA* control for unsynchronised and nocodazole treated samples. Positive ratio indicates enrichment in BidWT sample. P Value calculated via ANOVA from three independent replicates.

To validate mBid-BirA* fusions, we compared the pro-apoptotic function of truncated Bid (tBid) fused to either YFP or BirA*. Both induced similar levels of apoptosis when transiently expressed in HEK-293T cells (Fig.S1A). A venus-BirA* fusion did not induce apoptosis. We next validated tBid-BirA* dependent biotin labelling. Due to the potent pro-apoptotic activity of tBid, cell lines stably expressing tBid-BirA* could not be generated. Therefore, HEK-293T cells transiently expressing either tBid-BirA* or BirA* were grown in media with or without supplementary biotin for 18 hours. Whole cell lysates were probed with streptavidin or anti-myc (Fig.S1B). The expressed BirA*-fusion proteins both showed self-labelling in the presence of biotin. To determine if tBid-BirA* could biotinylate known binding partners, we transiently expressed GFP-Bcl-XL in HEK-293T cells, either alone or with tBid-BirA*. Transfected cells were supplemented with biotin, whole cell lysates (WCL) subjected to streptavidin affinity purification and blotted for GFP or biotin (Fig.S1C). GFP-Bcl-XL bound streptavidin beads only when co-expressed with tBid-BirA*, confirming BioID was able to capture interactions between tBid and anti-apoptotic Bcl-2 proteins *in situ*. Finally, we used immunofluorescence microscopy to visualise biotin in cells expressing either tBid-BirA* or BirA* (Fig.S1D). Only cells expressing either tBid-BirA* or BirA* were positive for biotin. Notably, tBid-BirA* expressing cells showed mitochondrial localisation of biotin.

In order to identify mitosis specific interacting partners, we used quantitative, label free mass-spectrometry with HeLa cells stably expressing full-length (FL) mBid-BirA* (Fig.1A), which allow robust enrichment in mitosis. We generated stable HeLa lines expressing mBidWT-BirA* or venus-BirA*. We also generated HeLa lines expressing Bid variants that were either not phosphorylated in mitosis (mBidS66A-BirA*), or where the BH3-domain required for interacting with other Bcl-2 proteins had been made non-functional (mBidG94E-BirA*). Fusion proteins were expressed from a lentivirus co-expressing tagBFP downstream of a T2A cleavage sequence, allowing FACS selection of stable lines with similar expression of each bait (Fig. 1B). Cells were maintained in biotin containing media, either in nocodazole or unsynchronised, for 16 hours, which allowed enrichment for labelling in mitosis vs. interphase (Fig. 1C). Labelled proteins were isolated by streptavidin affinity purification (AP), analysed by LC-MS/MS, and quantified using Progenesis QI. We performed three independent experiments, typically identifying >400 proteins within each sample (Fig.1D). Enrichment of individual proteins was determined by comparing their abundance in each mBid-BirA* sample to the experimentally paired venus-BirA* control. To prioritize proteins for further analysis, selection was restricted to those that were enriched in mBidWT-BirA* cells in all three experimental repeats (Tables 1 and 2).

**Table 1.** Summary of enrichment data in unsynchronised HeLa cells, from three independent experiments (R1, R2 and R3) comparing mBidWT-BirA* relative to venus-BirA*. Proteins are ordered by mean enrichment (Log2 ratio mBidWT-BirA* vs. venus-BirA*). Only proteins enriched in all three repeats are shown. Those highlighted in orange have CRAPome scores greater than 20%. VDAC2 is highlighted in red. Bid (highlighted in green) is the mouse protein used as bait.

| Gene Name | Description/Protein name | CRAPome % | Sequence coverage | Log2(ratio) |  |  |  |
| --- | --- | --- | --- | --- | --- | --- | --- |
|  |  |  |  | R1 | R2 | R3 | Mean |
| *BID* | BH3-interacting domain death agonis | 1.46 | N/A | 3.094 | 3.709 | 5.261 | 4.021 |
| *DDX3X* | ATP-dependent RNA helicase DDX3X | 51.58 | 25.0% | 0.595 | 1.327 | 1.894 | 1.272 |
| *HNRNPDL* | Heterogeneous nuclear ribonucleoprotein D-like | 33.82 | 6.3% | 0.362 | 0.411 | 2.660 | 1.144 |
| *EIF4G1* | Eukaryotic translation initiation factor 4 gamma 1 | 24.82 | 14.0% | 2.489 | 0.249 | 0.646 | 1.128 |
| *FLNC* | Filamin-C | 35.77 | 4.4% | 0.016 | 0.541 | 2.190 | 0.916 |
| *EIF4A1* | Eukaryotic initiation factor 4A-I | 46.23 | 45.0% | 0.262 | 1.085 | 1.327 | 0.891 |
| *CAND1* | Cullin-associated NEDD8-dissociated protein 1 | 21.41 | 6.3% | 0.157 | 1.904 | 0.550 | 0.870 |
| *XRN2* | 5'-3' exoribonuclease 2 | 23.60 | 2.9% | 0.568 | 0.379 | 1.413 | 0.787 |
| *SEPT9* | Septin 9, isoform CRA a | 19.22 | 35.0% | 0.309 | 1.482 | 0.448 | 0.746 |
| *SLC25A11* | Mitochondrial 2-oxoglutarate/malate carrier protein (Fragment) | 13.38 | 8.8% | 0.073 | 0.757 | 1.389 | 0.739 |
| *SON* | Protein SON | 17.27 | 0.7% | 0.069 | 0.382 | 1.564 | 0.672 |
| *RPA1* | Replication protein A 70 kDa DNA-binding subunit | 20.44 | 8.6% | 0.233 | 0.440 | 1.328 | 0.667 |
| *RPN1* | Dolichyl-diphosphooligosaccharide--protein glycosyltransferase subunit 1 | 22.14 | 18.0% | 1.114 | 0.428 | 0.374 | 0.639 |
| *DSP* | Desmoplakin | 30.66 | 17.0% | 0.508 | 0.772 | 0.537 | 0.606 |
| *NAT10* | RNA cytidine acetyltransferase | 17.03 | 4.5% | 0.091 | 0.640 | 1.053 | 0.595 |
| *VDAC2* | Voltage-dependent anion channel 2 | 14.11 | 49.0% | 0.193 | 1.361 | 0.143 | 0.566 |
| *RBBP6* | E3 ubiquitin-protein ligase RBBP6 | 9.49 | 1.9% | 0.583 | 0.313 | 0.686 | 0.527 |
| *HSPA1L* | Heat shock 70 kDa protein 1-like | 94.65 | 43.0% | 1.040 | 0.037 | 0.494 | 0.524 |
| *H2AFY* | Histone H2A | 16.55 | 5.9% | 0.770 | 0.304 | 0.378 | 0.484 |
| *SFPQ* | Splicing factor, proline- and glutamine-rich | 48.91 | 7.2% | 0.070 | 0.642 | 0.659 | 0.457 |
| *DYNC1H1* | Cytoplasmic dynein 1 heavy chain 1 | 31.87 | 3.1% | 0.081 | 0.388 | 0.873 | 0.447 |
| *RPL23A* | 60S ribosomal protein L23a (Fragment) | 54.50 | 20.0% | 0.091 | 0.608 | 0.596 | 0.432 |
| *MAP4* | Microtubule-associated protein | 20.44 | 11.0% | 0.345 | 0.767 | 0.165 | 0.426 |
| *STAT1* | Signal transducer and activator of transcription 1-alpha/beta | 5.60 | 12.0% | 0.056 | 0.231 | 0.825 | 0.371 |
| *HUWE1* | HECT, UBA and WWE domain containing 1 | 18.00 | 0.3% | 0.349 | 0.087 | 0.623 | 0.353 |
| *HSPA9* | Stress-70 protein, mitochondrial (Fragment) | 74.70 | 25.0% | 0.774 | 0.060 | 0.196 | 0.343 |
| *PCBP2* | Poly(rC)-binding protein 2 (Fragment) | 45.99 | 26.0% | 0.125 | 0.737 | 0.059 | 0.307 |
| *DLST* | Dihydrolipoamide S-succinyltransferase (E2 component of 2-oxo-glutarate complex) | 13.87 | 4.4% | 0.150 | 0.205 | 0.532 | 0.296 |
| *ABCD3* | ATP-binding cassette sub-family D member 3 | 9.73 | 3.5% | 0.003 | 0.281 | 0.567 | 0.284 |
| *DKFZp686L20222* | Putative uncharacterized protein DKFZp686L20222 | 19.71 | 26.0% | 0.265 | 0.289 | 0.291 | 0.282 |
| *CORO1B* | Coronin | 9.25 | 38.0% | 0.011 | 0.416 | 0.336 | 0.254 |
| *DNA helicase* | DNA helicase | N/A | 11.0% | 0.274 | 0.217 | 0.047 | 0.179 |
| *PAICS* | Multifunctional protein ADE2 | 25.79 | 40.0% | 0.101 | 0.121 | 0.268 | 0.164 |
| *PRDX1* | Peroxiredoxin-1 | 61.56 | 45.0% | 0.057 | 0.019 | 0.292 | 0.122 |
| *CSDE1* | Cold shock domain containing E1, RNA-binding | 16.30 | 21.0% | 0.064 | 0.241 | 0.021 | 0.109 |

**Table 2.** Summary of enrichment data in nocodazole treated HeLa cells, from three independent experiments (R1, R2 and R3) comparing mBidWT-BirA* relative to venus-BirA*. Proteins are ordered by mean enrichment (Log2 ratio mBidWT-BirA* vs. venus-BirA*). Only proteins enriched in all three repeats are shown. Those highlighted in orange have CRAPome scores greater than 20%. VDAC2 is highlighted in red. Bid (highlighted in green) is the mouse protein used as bait.

| Gene Name | Description/Protein name | CRAPome% | Sequence coverage | Log2(ratio) |  |  |  |
| --- | --- | --- | --- | --- | --- | --- | --- |
|  |  |  |  | R1 | R2 | R3 | Mean |
| *BID* | BH3-interacting domain death agonist (mouse) | 1.46 | N/A | 2.416 | 2.728 | 4.072 | 3.072 |
| *RPA1* | Replication protein A 70 kDa DNA-binding subunit | 20.44 | 2.1% | 0.179 | 0.228 | 4.966 | 1.791 |
| *G6PD* | Glucose-6-phosphate 1-dehydrogenase | 5.11 | 5.2% | 0.446 | 0.003 | 4.326 | 1.592 |
| *FH* | Fumarate hydratase isoform 1 (Fragment) | 5.84 | 8.0% | 0.326 | 1.219 | 2.902 | 1.482 |
| *BRAP* | BRCA1-associated protein | 4.62 | 2.2% | 0.462 | 0.242 | 3.442 | 1.382 |
| *DDX3X* | ATP-dependent RNA helicase DDX3X | 51.58 | 15.0% | 0.449 | 0.696 | 2.668 | 1.271 |
| *MYH14* | Myosin-14 | 32.12 | 37.0% | 1.025 | 0.099 | 2.656 | 1.260 |
| *CAPG* | Macrophage-capping protein | 1.22 | 16.0% | 1.056 | 0.061 | 2.014 | 1.044 |
| *NCAPG* | Non-SMC condensin I complex, subunit G | 14.36 | 2.8% | 0.822 | 0.094 | 2.107 | 1.007 |
| *VDAC2* | Voltage-dependent anion channel 2 | 14.11 | 35% | 0.812 | 0.914 | 1.154 | 0.960 |
| *LIMCH1* | LIM and calponin homology domains-containing protein 1 (Fragment) | 5.35 | 2.4% | 0.068 | 0.189 | 2.532 | 0.930 |
| *SLC25A31* | ADP/ATP translocase 4 | 42.82 | 37.0% | 0.091 | 0.288 | 2.392 | 0.924 |
| *EIF4G1* | Eukaryotic translation initiation factor 4 gamma 1 (Fragment) | 24.82 | 19.0% | 0.468 | 0.371 | 1.804 | 0.881 |
| *ARCN1* | Archain 1, isoform CRA_b | 22.38 | 9.0% | 0.096 | 0.208 | 2.311 | 0.872 |
| *NUP155* | Nucleoporin 155kDa, isoform CRA_a | 11.92 | 5.5% | 0.647 | 0.066 | 1.793 | 0.836 |
| *PCBP2* | Poly(rC)-binding protein 2 (Fragment) | 45.99 | 25.0% | 0.706 | 0.299 | 1.491 | 0.832 |
| *MCCC1* | Methylcrotonoyl-CoA carboxylase subunit alpha, mitochondrial | 21.41 | 15.0% | 0.006 | 0.113 | 2.367 | 0.829 |
| *HSPA9* | Stress-70 protein, mitochondrial (Fragment) | 74.70 | 26.0% | 1.143 | 0.040 | 1.143 | 0.776 |
| *HSPA8* | Heat shock cognate 71 kDa protein (Fragment) | 96.35 | 40.0% | 0.198 | 0.250 | 1.855 | 0.768 |
| *EIF4A1* | Eukaryotic initiation factor 4A-I (Fragment) | 46.23 | 44.0% | 0.250 | 0.842 | 0.852 | 0.648 |
| *DLST* | Dihydrolipoamide S-succinyltransferase (E2 component of 2-oxo-glutarate complex) | 13.87 | 4.4% | 0.138 | 0.326 | 1.414 | 0.626 |
| *CHD3* | Chromodomain-helicase-DNA-binding protein 3 | 13.63 | 2.7% | 1.232 | 0.324 | 0.146 | 0.567 |
| *HNRNPDL* | Heterogeneous nuclear ribonucleoprotein D-like | 33.82 | 6.3% | 0.511 | 0.327 | 0.674 | 0.504 |
| *AP3B1* | Adaptor-related protein complex 3 beta 1 subunit isoform 1 (Fragment) | 9.98 | 3.0% | 0.352 | 0.218 | 0.849 | 0.473 |
| *EPRS* | Bifunctional glutamate/proline--tRNA ligase | 38.20 | 25.0% | 0.668 | 0.145 | 0.568 | 0.460 |
| *EPS15L1* | Epidermal growth factor receptor substrate 15-like 1 | 6.57 | 15.0% | 0.402 | 0.149 | 0.830 | 0.460 |
| *CHD5* | Chromodomain-helicase-DNA-binding protein 5 | 14.11 | 1.3% | 0.129 | 0.095 | 1.137 | 0.454 |
| *AHNAK2* | Protein AHNAK2 | 4.87 | 9.5% | 0.339 | 0.290 | 0.475 | 0.368 |
| *HUWE1* | HECT, UBA and WWE domain containing 1 | 18.00 | 0.3% | 0.182 | 0.049 | 0.737 | 0.323 |
| *DSP* | Desmoplakin | 30.66 | 21.0% | 0.132 | 0.363 | 0.272 | 0.256 |
| *PABPC3* | Polyadenylate-binding protein 3 | 31.63 | 8.6% | 0.121 | 0.309 | 0.270 | 0.233 |
| *ABCF3* | ATP-binding cassette sub-family F member 3 isoform 1 (Fragment) | 0.73 | 5.9% | 0.143 | 0.209 | 0.111 | 0.154 |
| *PDLIM1* | PDZ and LIM domain protein 1 | 7.79 | 35.0% | 0.029 | 0.185 | 0.095 | 0.103 |

Surprisingly, proteins enriched in both unsynchronised and mitotic mBid-BirA* cells did not include any associated with Bid’s documented roles within the intrinsic and extrinsic apoptotic pathways (Fig.2A). The STRING database (31, 32) predicted that the top 10 functional partners of Bid would be associated with these pathways, including other Bcl-2 proteins, components of the death induced signaling complex (DISC) and caspase proteases (Fig.2B). However, the proteins enriched by mBid-BirA* did not overlap with this predicted network. Curious if the STRING-predicted interaction partners were detected in any of the experimental replicates, we lowered the stringency to include proteins enriched in one or two experimental repeats (Fig.2C). Bax was enriched with mBidWT-BirA* in two repeats, while Bcl-XL was identified in one. However, for both proteins fewer than 3 peptides were detected, making any quantitative comparison unreliable. Having shown above that tBid-BirA* was capable of biotinylating Bcl-XL in cells (Fig.S1C), the inefficient labelling of Bcl-XL and Bax suggested that these proteins were not predominantly in close proximity to full-length mBid-BirA* in non-apoptotic cells.

**Figure 2.**
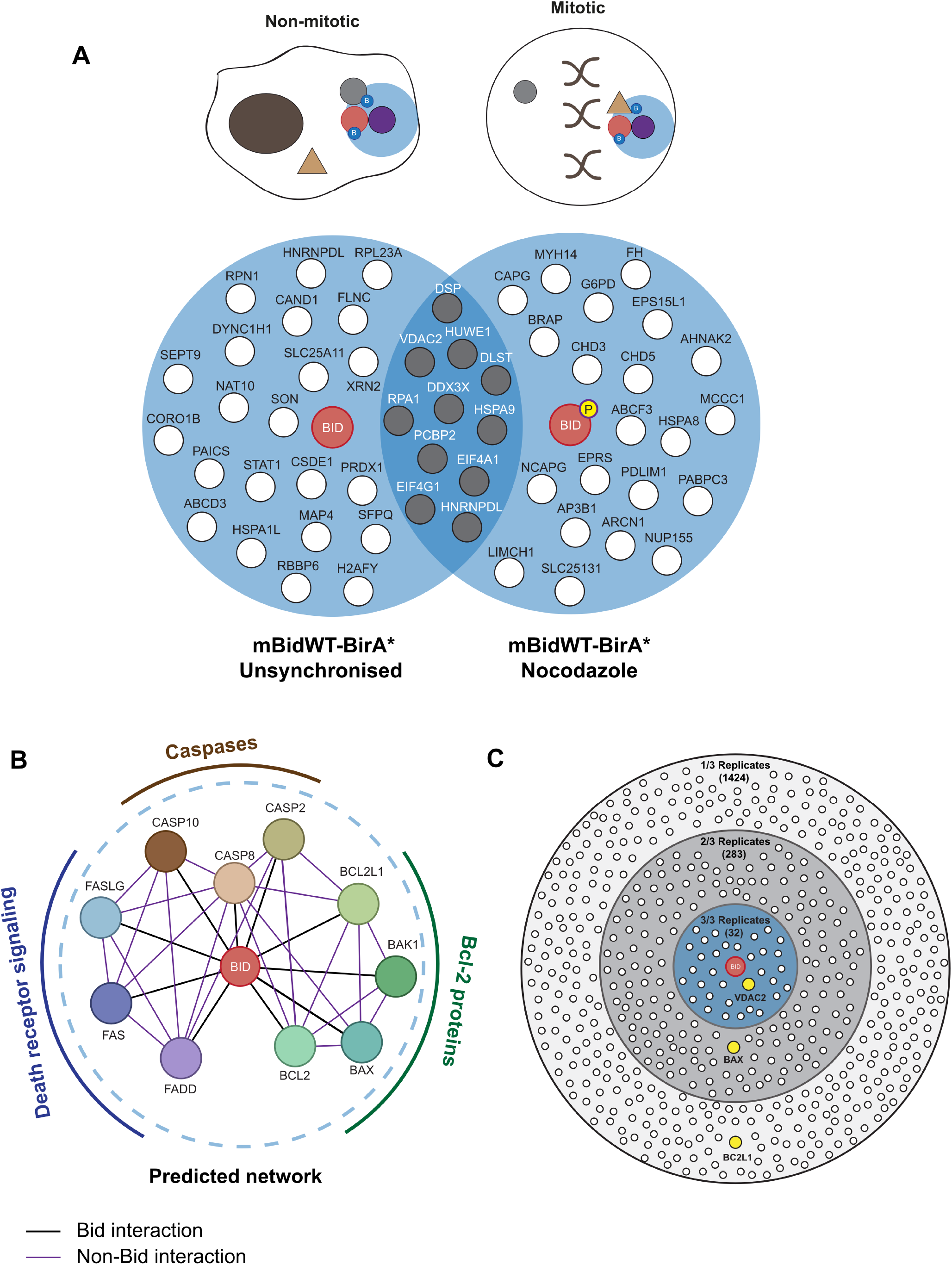
FL-Bid based BioID mass spectrometry screen does not enrich canonical Bcl-2 family proteins. A. Venn diagram showing proteins identified as enriched by mBidWT-BirA* vs. venus-BirA* under unsynchronised and nocodazole treated conditions. Only proteins identified as enriched in all three independent experiments are shown. Proteins identified in both conditions are shaded grey. There is divergence between the predicted interaction network based on previous apoptosis focused literature shown in (B), and the proteins identified by mBidWT-BirA* screen under non-apoptotic conditions. B. The predicted protein interaction network for Bid derived from top 10 highest scoring predicted functional partners according to STRING (experimentally validated interactions and interactions in curated databases only). Nodes in blue represent death receptors/death receptor ligands, nodes in brown represent caspases, while nodes in green represent Bcl-2 proteins. C. Bcl-2 family proteins are identified in the mBid-BirA* data set when the stringency is lowered. 32 proteins, including VDAC2, were identified as being enriched in mBid-BirA* vs. venus-BirA* in 3 out of 3 replicates (blue inner circle). 283 proteins were enriched in 2 out of 3 experimental replicates, including Bax. 1424 proteins were identified in 1 out of 3 experimental replicates, including Bcl-XL (BCL2L1).

Together, these data suggested that Bcl-2 family proteins may not represent the main interaction partners for full-length Bid in live cells.

### VDAC2 is a potential Bid partner in mitosis

A common issue with all AP-MS experiments is the detection of significant background contaminants. Given that we did not detect predicted Bid interacting partners, we were concerned that BioID might have predominantly isolated non-specific proteins. It was, therefore, important to validate candidate proteins. We first compared the sequence coverage of each protein and their relative representation in the contaminant repository for affinity purification (CRAPome), which collates proteins commonly found within negative control AP-MS datasets (Fig.3A; Tables 1 and 2) (33). Falling within the criteria of high sequence coverage and low representation in the CRAPome was the mitochondrial porin, voltage-dependent anion-selective channel protein 2 (VDAC2), which has been previously linked to the function of multiple Bcl-2 family proteins (34–37). Other potential candidate proteins included the E3-ubiquitin ligase Huwe1 (MULE/ARFBP1), known to regulate the levels of Mcl-1, and glucose-6-phosphate dehydrogenase (G6PD), which is a target of PLK1 in mitosis (38, 39).

**Figure 3.**
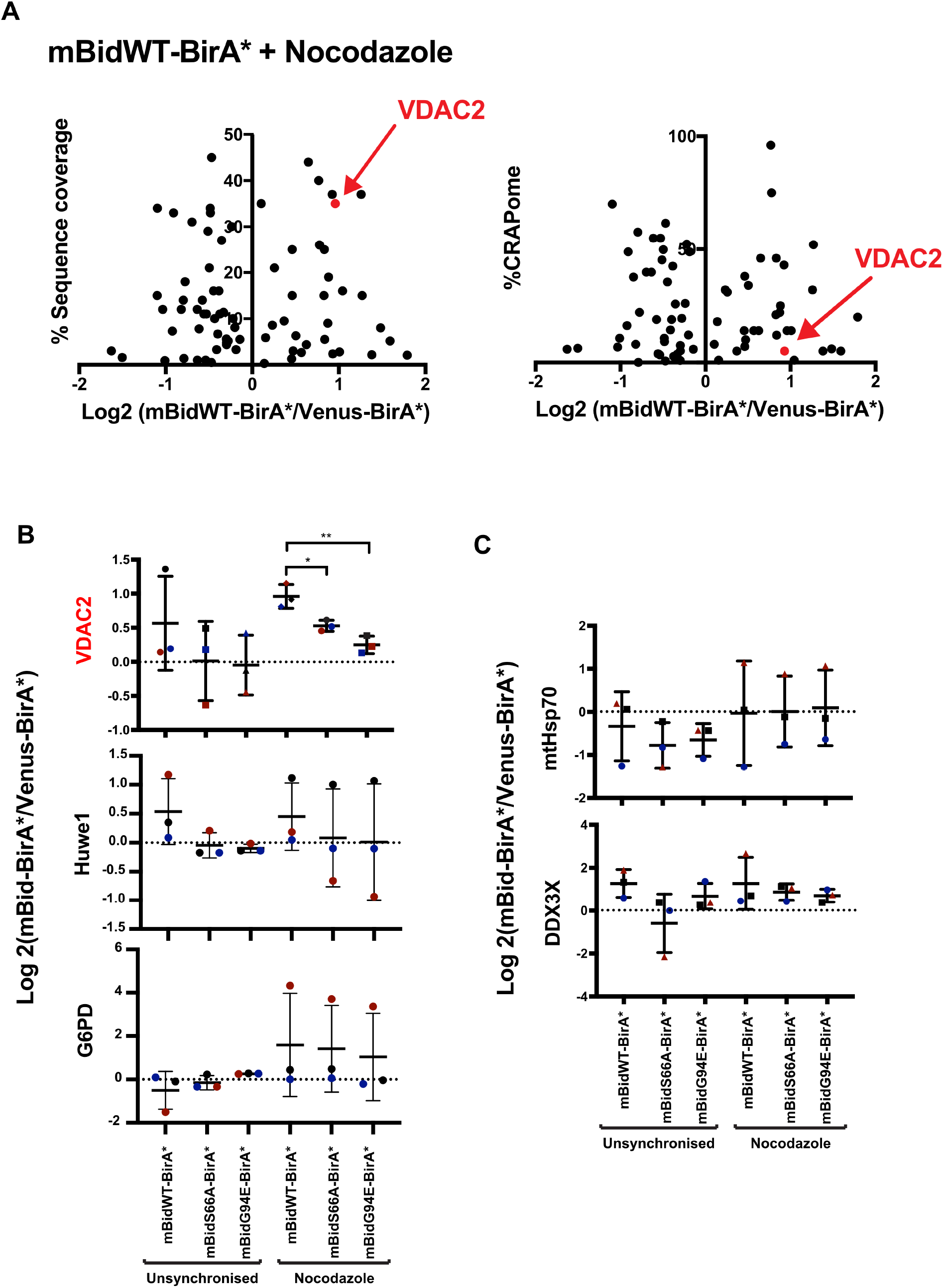
BioID identifies the voltage dependent anion channel 2, VDAC2, as showing Bid phosphorylation dependent enrichment in mitotic cells. A. Scatter plots comparing mean fold enrichment for mBidWT-BirA* vs. venus-BirA* with sequence coverage (left panel) and abundance in published negative control AP-MS data sets (%CRAPome, right panel). VDAC2 is indicated in both plots as having high sequence coverage, and in low percentage in CRAPome. B. Summary of MS analysis from showing protein abundance fold change for VDAC2, the HECT-domain containing E3-ubiquitin ligase Huwe1, and glucose-6-phosphate dehydrogenase (G6PD). Data represents mean and SD of 3 independent replicates. Significance calculated by unpaired t Tests, * P<0.05, ** P<0.01. C. Summary of MS analysis from showing protein abundance fold change for mtHSP70 and DDX3X, both of which show relatively high scores for their relative frequency in control AP-MS data.

To gain further confidence for which candidates might associate with Bid specifically in mitosis, we compared their enrichment in unsynchronised and mitotic cells expressing either mBidWT-BirA*, mBidS66A-BirA* or mBidG94E-BirA* (Fig.3B). As neither mBidS66A nor mBidG94E increased priming of mitotic colon carcinoma cells (21), we predicted that a specific regulatory partner would show differential enrichment with these baits compared to mBidWT. Of the potential candidates, VDAC2 was only enriched in mitotic cells expressing the mBidWT-BirA* bait compared to unsynchronised cells. Furthermore, VDAC2 enrichment was significantly less in mitotic cells expressing either BidS66A-BirA* or BidG94E-BirA*. Neither Huwe1 nor G6PD showed specific enrichment associated with mitosis (Fig.3B). Huwe1 showed a slight enrichment with mBidWT-BirA*. Proteins identified that are commonly found in control AP-MS data sets showed no specific enrichment in mitosis or with mBid variants. For example, both mitochondrial heat shock protein 70 (*HSPA9*, mtHsp70) and ATP dependent RNA helicase (*DDX3X,* DDX3X) showed no difference between lines expressing mBidWT-BirA* or the S66A and G94E variants (Fig.3C). Notably, VDAC1, which is ~10-fold more abundant than VDAC2 in HeLa cells (40), was not enriched in any samples. Importantly, all quantification was performed using only non-conflicting peptides, so sequence similarity between different VDAC isoforms was not a confounding issue.

Thus, BioID identified VDAC2 as the principal candidate for a Bid phosphorylation specific role in mitotic cells.

### VDAC2 couples Bid phosphorylation with increased apoptotic priming in mitosis

To determine if VDAC2 influenced apoptotic priming in mitosis, we employed a knock CRISPR/Cas9 out (ko) strategy in cells that showed Bid phosphorylation dependent priming in mitosis. We compared the human mammary carcinoma cell lines, MCF-7 and MDA-MB-231, for susceptibility to apoptosis *vs*. slippage following treatment with Taxol. Single cell fate profiles were obtained through live imaging of unsynchronised cells, in the presence or absence of Taxol, over 65 hours, quantifying entry into mitosis, normal division, apoptosis and slippage (Fig.S2). Few untreated cells underwent apoptosis. However, in Taxol MCF-7 cells predominantly underwent apoptosis, whereas MDA-MD-231 cells were prone to slippage (Fig.S2D). This inter- and intra-line variation in apoptosis is consistent with previous studies (3, 41).

We chose MCF7 cells for generating a VDAC ko line, as these predominantly underwent death in mitosis. We first confirmed whether Bid phosphorylation set MCF7 apoptotic priming in mitosis by stably knocking down endogenous hBid using a lentiviral shRNA (shBid). We simultaneously expressed mBid-GFP variants, using an ubiquitin promoter to achieve close to endogenous expression levels (21). MCF-7 lines were generated expressing either shBid alone, or with mBidWT-GFP, mBidS66A-GFP or mBidG94E-GFP, using FACS to select similar expression for each variant (Fig. 4A). Immunoblotting for each variant in unsynchronised and mitotic cells indicated that mBidWT-GFP and mBidG94E-GFP were phosphorylated in mitosis, whereas mBidS66A-GFP was not (Fig.4A). Near infrared immunoblotting was used to quantify hBid knockdown (Fig.S3A).

**Figure 4.**
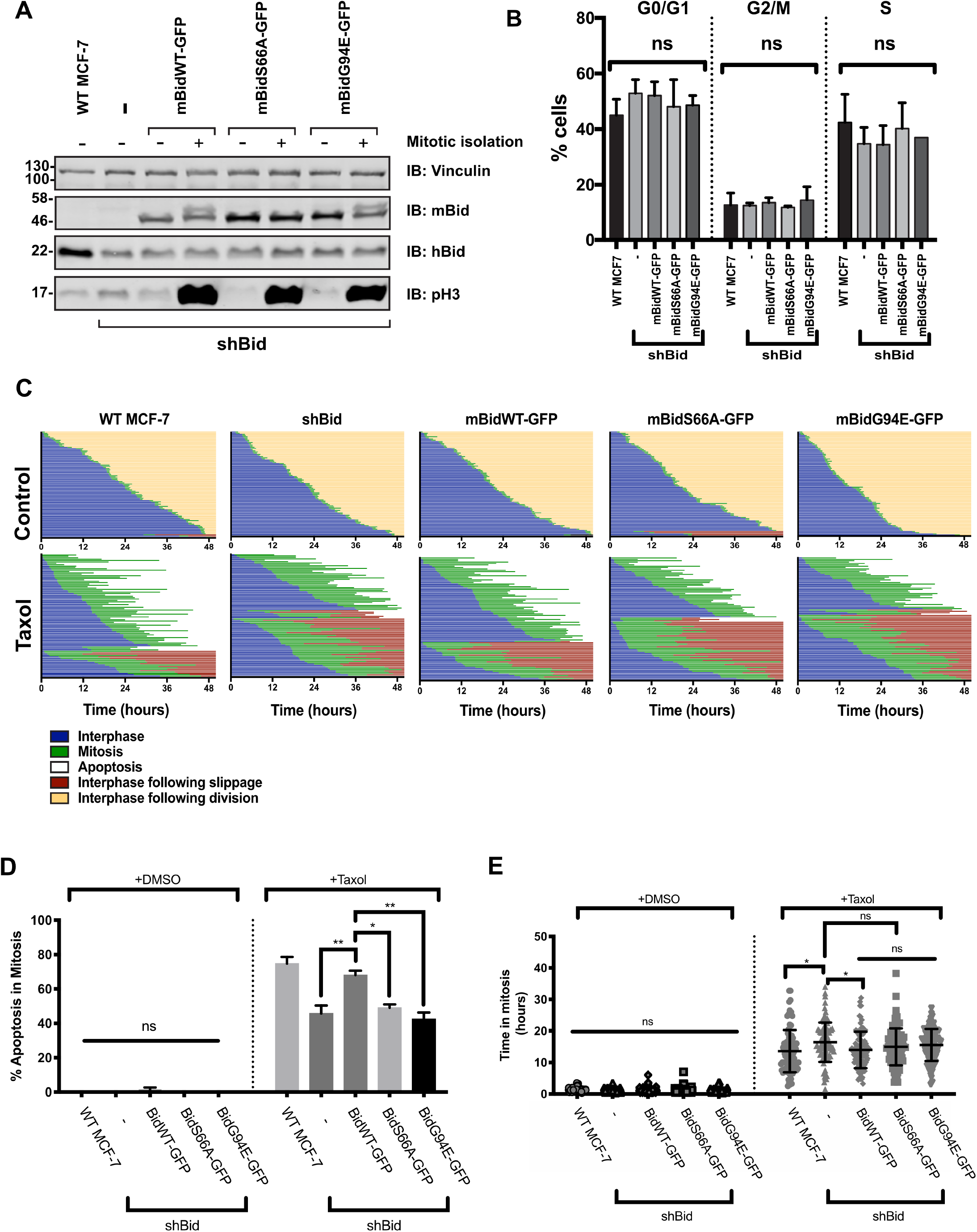
Phosphorylation of Bid regulates apoptotic priming in mitotic breast cancer cells. A. MCF-7 cell lines were generated stably expressing either shBid alone or with the indicated mouse Bid-GFP variant (mBidWT-GFP, mBidS66A-GFP, mBidG94E-GFP). Cells were either untreated (−) or enriched for mitosis with 18 hour treatment in nocodazole followed by shake off (+). Whole cell lysates were immunoblotted with the indicated antibodies. Anti-vinculin serves as loading control, while anti-phospho-Histone H3 (pH3) indicates mitosis. B. Cell cycle distribution for the MCF7 lines in A, quantified through DAPI staining and flow cytometry. Data were obtained using unsynchronized cells. Data represents mean and SD of 2 independent repeats. C. Single cell fate profiles of the MCF-7 lines in A, untreated control or treated with 1μM Taxol over 50 hours. Data represents 90 cells tracked over 3 independent experiments. D. Summary of apoptosis in mitosis for the data in C for the indicated MCF7 cell lines. Mean and SD are shown. Data were analysed by one-way ANOVA, followed by Tukey’s multiple comparison test (ns = non-significant; *** = p<0.001; **** = p<0.0001). E. Summary of duration of mitosis, all fates included, for the data in C. Mean and SD plotted. Data analysed by one-way ANOVA, followed by Tukey’s multiple comparison test (ns = non-significant; *** = p<0.001; **** = p<0.0001).

Flow cytometry showed that Bid expression, phosphorylation or apoptotic function had no effect on cell cycle progression (Fig.4B). We compared single cell fates for parental MCF-7 (WT MCF-7) cells with those expressing shBid, either alone or with each mBid-GFP variant. In the absence of Taxol there was no significant apoptosis or slippage in any of the lines (Fig4C,D). However, in the presence of Taxol, Bid knockdown significantly shifted MCF-7 cell fate toward slippage (Fig.4C,D). This was corrected by expression of mBidWT-GFP, but not by mBidS66A-GFP or mBidG94E-GFP. Knock down of Bid or expression of any of the FL Bid variants had no impact on the time untreated cells spent in mitosis (Fig.4C,E). Taxol clearly increased the time all cells spent in mitosis. Loss of Bid expression slightly increased the time in Taxol arrested mitosis relative to parental cells, which we suspect is accounted for by the reduced apoptosis. Thus, Bid phosphorylation and pro-apoptotic function were required for increasing priming in MCF-7 cells during mitosis. The level of apoptotic priming in all the shBid MCF7 lines could be reset to that of WT MCF7 cells by treating with the BH3-mimetic, ABT737 (Fig.S3). ABT737 alone had no effect. MDA-MB-231 cells stably expressing shBid with either mBidWT-GFP, mBidS66A-GFP or mBidG94E-GFP were also examined (Fig.S3D). As with MCF-7 cells, the Bid variants by themselves did not alter normal mitosis (Fig.S3E). Interestingly, although the baseline for priming in MDA-MB-231 was higher than for MCF-7, there was still an observable a reduction in apoptosis in Taxol for cells expressing Bid-S66A-GFP or BidG94E-GFP compared with mBidWT-GFP.

MCF7 cells showed Bid phosphorylation dependent priming in mitosis. We therefore targeted the start of the protein-coding region of the *VDAC2* gene in MCF7 cells using CRISPR/Cas9. Three independent clones containing indels were identified, termed D11, D6 and D4 (Fig.5A). MCF-7 cells have three copies of *VDAC2*. Sequencing indicated that for D11, all three copies contained frame-shift insertions. D6 had two frame shift insertions and one WT allele, and D4 two frame shift insertions and one allele with a 3 bp deletion just after the start of the coding sequence.

**Figure 5.**
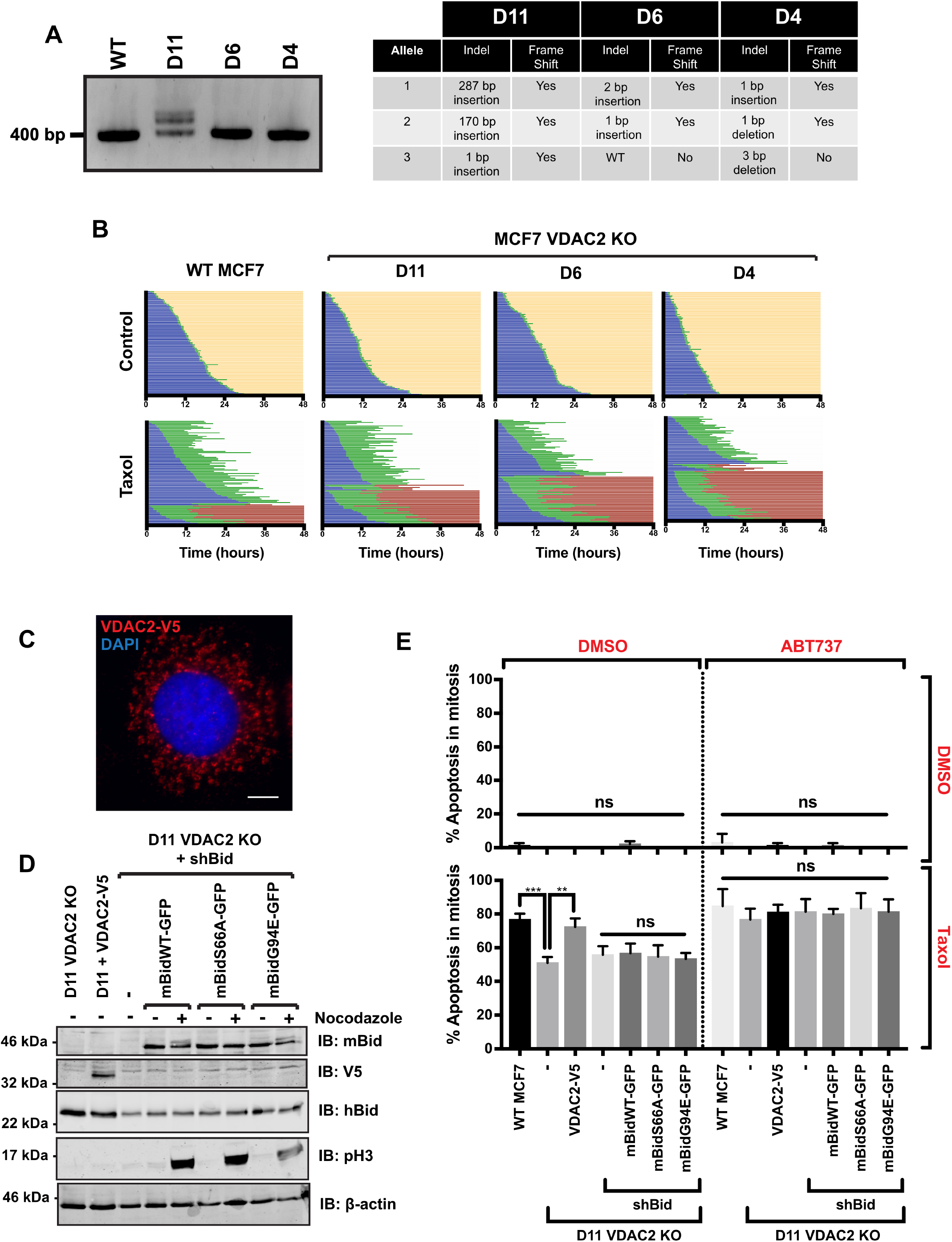
VDAC2 coordinates Bid phosphorylation dependent apoptotic priming in mitosis. A. Single MCF-7 cell clones were grown following CRIPSR/Cas9 targeting of *VDAC2*. Left hand panel - agarose gel electrophoresis of 300 bp PCR product at *VDAC2* CRISPR target PAM site. Right hand panel - table displaying indels caused by CRISPR/Cas9 determined by direct DNA sequencing of the amplified regions. B. Single cell fate profiles of the three MCF-7 *VDAC2* KO lines in (A), untreated (control) or treated with 1μM Taxol over 48 hours. Data represents 90 cells over 3 independent experiments. C. D11 *VDAC2* KO cells stably expressing VDAC2-V5 were fixed and immunostained for V5. A T2A-tagRFP sequence was included downstream of VDAC2-V5 coding sequence to allow selection by FACS. Scale bar = 10μm. D. D11 *VDAC2* KO MCF-7 cell lines were generated stably expressing VDAC2 - V5, shBid alone or shBid in conjunction with the indicated mouse Bid-GFP variant (mBidWT-GFP, mBidS66A-GFP, mBidG94E-GFP). Cells were either untreated or enriched for mitosis with 18 hour treatment in nocodazole followed by shake off. Whole cell lysates were immunoblotted with the indicated antibodies. β-actin serves as loading control, while phospho-Histone H3 (pH3) indicates mitosis. E. Summary of apoptosis in mitosis for single cell fate analysis of lines in D, untreated or treated with 1μM Taxol and/or 5 μM ABT-737 over 48 hours (single cell traces in Figure S6). Data represents 90 cells tracked over 3 independent experiments. Error bars represent SD Data analysed by one-way ANOVA, followed by Tukey’s multiple comparison test. ns = non-significant, ** = p <0.01, *** = p <0.001.

We performed cell fate analysis to compare WT MCF-7 cells to the three VDAC2 ko lines (Fig.6B). There were no differences in normal mitotic progression in untreated cells. However, when mitosis was prolonged with Taxol, all three VDAC2 ko lines showed a shift toward slippage compared to WT cells (Fig.5B). Furthermore, the proportion of each VDAC2 ko line undergoing slippage was comparable to that seen with shBid MCF-7 cells, or those expressing mBidS66A or mBidG94E (cf. Fig.4C). Re-expression VDAC2 tagged with a C-terminal V5 epitope (VDAC-V5; Fig.5C,D) in the MCF-7 D11 line restored the level of apoptosis in Taxol to that of WT MCF-7 cells (Fig.5E).

We asked if VDAC2 was required for coupling Bid phosphorylation to mitotic priming. The D11 line was therefore stably transduced with shBid alone, or shBid along with mBidWT-GFP, mBidS66A-GFP or mBidG94E-GFP (Fig.5D). Notably, both mBidWT-GFP and mBidG94E-GFP were still phosphorylated in mitotic D11 cells, seen with the mobility shift in the presence of Nocadazole. Cell fate profiling in the presence or absence of Taxol compared each mBid-GFP D11 line to WT MCF7, the D11 ko and D11-VDAC2-V5 cells (Fig.S4). Bid knockdown had no further effect on the level of apoptosis beyond that already brought about by *VDAC2* deletion (Fig.5E). mBidWT-GFP, mBidS66A-GFP or mBidG94E-GFP expressing D11 cells all showed the same proportion of death vs. slippage in Taxol as the D11 VDAC2 ko (Fig.5E). Importantly, the level of death in D11 cells was comparable to shBid MCF-7 cells (cf. Fig.2, Fig.S2). Apoptosis in all lines was restored to the level of WT MCF-7 cells by treating with ABT-737 along with Taxol. Again, ABT-737 by itself did not induce apoptosis in any of the lines. These data show that VDAC2 is required to coordinate the changes in apoptotic priming in response to Bid phosphorylation in mitosis.

## Discussion

Cells constantly balance a multitude of pro-survival and pro-death signals. However, despite the current mechanistic understanding of apoptosis, it is still unclear how Bcl-2 proteins dynamically adjust this balance in response to rapidly changing signals. Using an unbiased proteomic approach to identify Bid vicinal proteins, we identified VDAC2 as a coordinator of dynamic apoptotic priming in mitotic cells. Somewhat surprisingly, we did not identify other Bcl-2 family proteins as the primary interactions with FL-Bid in non-apoptotic cells. The BioID proximity labelling approach was chosen to as proteins are biotinylated *in situ* and can then be isolated under stringent conditions. This allows isolation of integral membrane components without having to maintain direct protein-protein interactions. Furthermore, Bcl-2 proteins have a tendency to interact non-specifically when isolated in with detergents, complicating co-affinity purification approaches (29).

The lack of canonical Bcl-2 family proteins identified by BioID using FL-Bid, both in unsynchronized cells and those in mitosis, was unexpected. Bid was originally identified as an interacting partner of both Bcl-2 and Bax using a phage expression library and co-immunoprecipitation using non-ionic detergents (42). Subsequently, Bid was studied almost exclusively in the context of its cleavage by caspase 8, which converts it into a potent direct activator BH3-only protein (43, 44). Thus, interacting partners of Bid defined in the literature are those associated with its cleaved form in dead or dying cells. tBid-BirA* did label GFP-Bcl-XL in transiently expressing cells. Furthermore, a limited number of Bax and Bcl-XL peptides were detected in the AP-MS. Together, these data indicate that mBid-BirA* does label other Bcl-2 proteins. However, these interactions may occur principally in those cells that are undergoing apoptosis. It has been shown that FL-Bid is pro-apoptotic in the absence of caspase cleavage (45, 46). In non-apoptotic cells, FL-Bid may associate with regulatory proteins outside the core Bcl-2 family and established components of the extrinsic pathway. Other non-canonical interactions with BH3-only proteins, including Bid and Bad, have been identified (47, 48).

All AP-MS approaches suffer non-specific hits, commonly due to abundant proteins that are not sufficiently removed by washing, or sticky proteins that associate non-specifically with the beads or bait. A number of points supported VDAC2 as a genuine hit. Firstly, of the proteins we identified, VDAC2 was the only one enriched in mitotic cells only when Bid could be phosphorylated on S66 and had a functional BH3-domain. Thus, using mBidS66A-BirA* and mBidG94E-BirA*, as negative controls mitigated against non-specific interactions. Thus, VDAC2 enrichment showed a specificity for bait proteins that matched their function. Second, the three mammalian isoforms of VDAC show high sequence similarity, but have distinct roles (49). Importantly, although VDAC1 is 10-fold more abundant than VDAC2 in HeLa cells (40), it was not enriched by any of the mBid-BirA* variants. Thirdly, the functional validation through VDAC2 deletion, performed in a distinct cell line (MCF-7) from that used in the AP-MS screen, specifically uncoupled mitotic priming from Bid phosphorylation. Interestingly, Bid was still phosphorylated in mitotic VDAC2 ko cells, indicating that VDAC2 function lies downstream of potential Bid kinases. Loss of VDAC2 dampened priming in mitosis to the same extent as seen with Bid knock down, and their combined loss was not additive, showing that effect of Bid phosphorylation was mediated by VDAC2. Importantly, loss of VDAC2 did not simply make the cells resistant to apoptosis. The effect on apoptotic priming in mitosis due to the loss of either VDAC2, FL-Bid or both could be rescued by the BH3-mimetic ABT737. Taken together, these data indicate that VDAC2 coordinates FL-Bid functioning as a sensitizer BH3-only protein in this context.

A number of studies have linked VDAC2 to other Bcl-2 proteins, although the conclusions are often conflicting. Chemical cross-linking *in situ* identified VDAC2 with Bak, suppressing its activation (34). Conversely, other studies found VDAC2 functioned in Bak mitochondrial targeting, being required for tBid to activate Bak (35, 50). A genome-wide CRISPR/Cas9 screen in Mcl-1 deficient MEFs recently identified VDAC2 as a mediator of Bax, but not Bak, dependent apoptosis (37). Yet another study found that Bax retrotranslocation from mitochondria required VDAC2, and suggested that it functioned as a platform to coordinate the interaction between Bax and Bcl-XL (36). While the true role of VDAC2 is yet to be fully elucidated, VDAC2 is emerging as a nexus for Bcl-2 protein co-ordination on mitochondria. Our data here would support such a pivotal role in determining cellular response to apoptotic stimuli.

Finally, these results add to our understanding of how cells coordinate the dynamic changes in priming during mitosis. Mitosis represents a clinically important therapeutic target for drugs like Taxol, and numerous studies have implicated roles for different Bcl-2 family. A genome-wide screen in HeLa cells identified Bax as a key protein for Taxol mediated cell death, whereas the same approach in RKO cells did not directly identify any Bcl-2 proteins, but instead identified Myc dependent transcription of Bid, BIM and Noxa (22, 23). An siRNA approach in HeLa cells specifically targeted at Bcl-2 proteins identified roles for Noxa, Bim and Mcl-1 (51). Together, these studies indicate that the outcome of delayed mitotic exit is not determined by a single BH3-protein or multi-domain effector, but rather the complex interplay of many, the balance of which varies between and probably within cell types. Several Bcl-2 proteins undergo mitosis-specific post-translational modifications, including Bcl-2, Bcl-XL, Mcl-1, Bid and BimEL (18–21). Individually, these post translational modifications appear to have different roles. Phosphorylation of Mcl-1 promotes its degradation (24), whilst phosphorylation Bcl-XL and Bcl-2 increase their anti-apoptotic potency (18). However, collectively they result in the dynamic accumulation of apoptotic priming if anaphase is delayed (10). Our data now show that, unlike caspase 8 cleavage, phosphorylation of FL-Bid does not convert it into an activator BH3-only protein. Instead, phosphorylated FL-Bid functions more like a sensitiser, reversibly increasing apoptotic priming in mitosis. It is interesting to speculate that, in addition to its established function as a direct activator downstream of death receptors, a significant role for Bid may be as a sensitiser during mitosis.

Together, our results reveal Bid mediated mitotic priming is arbitrated *via* interactions outside of the core Bcl-2 family, highlighting how dynamic regulation of apoptotic priming is more complex that traditionally perceived. The data demonstrate thay BioID-based proteomics will allow dissection of the wider Bcl-2 family interactome in live cells. With the advent of Bcl-2 protein targeted drugs, it is essential to examine the wider interactions of these proteins in order to understand the complex layers of regulation which modulate their function.

## Methods

### Cell culture

MCF-7, BT-474, MDA-MB-231, SK-BR-3, MCF-10A, HEK-293T, and HeLa cells were acquired from ATCC (American Type Culture Collection). MCF-10A cells were grown in DMEM-F12 supplemented with 5% horse serum (v/v), 100 ng/ml cholera toxin, 20 ng/ml epidermal growth factor, 0.5 g/ml hydrocortisone, 10 mg/ml insulin, 100 U/ml penicillin and 100 mg/ml streptomycin. HeLa cells were grown in EMEM supplemented with 10% foetal bovine serum (v/v), 100 U/ml penicillin and 100 mg/ml streptomycin. All other cell lines were cultured in DMEM supplemented with 10% foetal bovine serum (v/v), 100 U/ml penicillin and 100 mg/ml streptomycin.

### Single cell fate tracking

1 μM Taxol (Sigma-Aldrich), 5 μM ABT-737 (Sigma-Aldrich) or DMSO diluted in complete growth media was added to cells 30 minutes prior to imaging. Images were acquired at 15 minute intervals for 48-60 hours on an AS MDW live cell imaging system (Leica) in bright-field using a 20x objective and utilising imaging software Image Pro 6.3 (Media Cybernetics Ltd). Point visiting was used to allow multiple positions to be imaged within the same time course and cells were maintained at 37°C and 5% CO_2_. Image stacks were analysed in ImageJ (National Institutes of Health), where single cells were identified and followed manually using cellular morphology to determine cell cycle progression and fate. Cells were described as being in mitosis from the initial point at which a cell began to round, till the first frame of cytokinesis or membrane blebbing. Cells that exhibited morphology associated with apoptosis such as membrane blebbing, shrinking of cell body, and in the later stages the formation of apoptotic bodies or secondary necrosis were described as having undergone apoptosis in mitosis. Cells that exited mitosis as a single cell, or as two or more non-identical daughter cells were grouped together under the definition of having undergone mitotic slippage.

### Antibodies, immunofluorescnce and immunoblotting

The following antibodies were used for immunoblotting and immunoflourescence: rabbit anti-Bid (ProteinTech, #10988-1-AP); mouse anti-Myc (Millipore, #05-724); rabbit anti-V5 (Sigma-Aldrich, # V8157); mouse anti-mtHsp70 (Thermo, # MA3-028); rabbit anti-GFP (Invitrogen, # PA1-9533); goat anti-β-actin (Abcam, # Ab8229); rabbit anti-phospho-Histone serine10 (Millepore, #06-570); mouse anti-vinculin (Abcam, # Ab9532). To detect biotinylated proteins we used DyLight 549 Streptavidin (Vector laboratories, # SA-5549) and IRDye680 Strepavidin (LiCor, #925-68079).

Immunofluorescent staining of cells was performed as previously described (7), using Alexafluor 594 and 488 conjugated secondary antibodies (Stratech). Immunostained cells were visualised using a Zeiss AxioImager M2 florescence microscope with 63x (NA 1.4) PLAN-APOCHROMAT objective. Images were acquired with Micro-Manager software and processed using ImageJ.

For immunoblotting, proteins were separated by SDS-PAGE and transferred to nitrocellulose. Following incubation with primary antibodies, proteins were detected using IrDye 800 and 680 secondary antibodies and an Odyssey CLx imager (LiCor). Quantification of blots was performed using ImageStudio (LiCor).

### Mitotic enrichment of cultured cells

Asynchronous cell populations were treated with nocodazole (Sigma-Aldrich, 200 ng/mL in complete growth media) for 16 hours to arrest cell in mitosis. Mitotic cells were then dislodged from the cell culture dish by tapping the vessel laterally against the bench. Media containing dislodged mitotic cells was removed and centrifuged at 350 RCF for 5 minutes to pellet cells ready for subsequent analysis.

### Flow cytometry analysis of cell cycle

Cell cycle distribution within a population was quantified by DNA content. Cycling cells were fixed with 70% ethanol and stained with DAPI. Stained cell samples were analysed on a BD Biosciences Fortessa flow cytometer, using Diva Version 8 (BD Biosciences) to collect raw data. Raw data was then analysed using Modfit LT (Version 5, Verity Software House) to quantify distribution between cell cycle phases.

### BioID mediated proximity labeling

For small scale assays such as Western blotting or immunofluorescent staining, cells expressing BirA* fusion proteins were seeded in a 6 well plate to approximately 80% confluence on day of labelling. Growth media was replaced with complete growth media supplemented with 50 μM biotin (Sigma-Aldrich), and cells cultured for a further 16 hours. Following incubation, biotin supplemented media was removed and replaced with regular growth media for 1 hour. Cells were then washed three times in PBS and assayed as appropriate.

For mass spectrometry experiments, cells expressing BioID fusion proteins were seeded into two 100 mm diameter dishes to be approximately 80% confluent next day. On the day of labelling, growth media was replaced with complete growth media supplemented with 50 μM biotin (Sigma-Aldrich) and cultured for a further for 16 hours, along with any other drug treatments. Following incubation, biotin supplemented media was removed and replaced with regular growth media for 1 hour to allow free biotin to diffuse out of cells. Cells were then washed three times in PBS and lysed in RIPA supplemented with 1x protease inhibitor cocktail (Millipore), 100 mM Na_3_VO_4_ and 500 mM NaF. Following lysis, lysates from both dishes were pooled, protein concentration measured and biotinylated proteins isolated via streptavidin-bead purification.

### Streptavidin affinity purification

Biotinylated proteins were isolated using MagReSyn^®^ streptavidin (MagReSyn^®^). 100 μL of bead slurry was decanted into a microcentrifuge tube and exposed to a magnetic field to separate the liquid and bead phases. The liquid phase was discarded and beads were washed three times in ice-cold lysis buffer. Appropriate volumes of cell lysate were added to beads to yield 1 mg of total protein, and the volume made up to a total of 1 mL with lysis buffer. Lysates and beads were incubated overnight at 4°C on an end-over-end tumbler. The next day beads were collected, washed twice in lysis buffer, once in urea wash buffer (2 M urea, 10 mM Tris-HCl pH 8.0), then three more times with lysis buffer. Following the final wash, beads were resuspended in 100 μL 1x lithium dodecyl sulphate sample buffer (Novex NuPAGE) supplemented with 2 mM biotin and 50 mM dithiothreitol (DTT) and incubated at 95°C for 5 minutes to elute bound proteins. Following elution, beads were collected and concentrated to 50 μL in a vacuum concentrator. 45 μL of sample was immediately subjected to SDS PAGE and processed for digestion in preparation for mass spectrometry, while the remaining 5 μL was analysed *via* western blotting to confirm biotinylation and protein enrichment.

### Mass spectrometry and protein identification

Affinity purified biotinylated proteins samples were briefly separated by SDS PAGE, then fixed and stained using InstantBlue (Expedeon). Gel fragments containing protein samples were digested with trypsin and peptides extracted under standard conditions ^40^. Peptides were subjected to liquid-chromatography tandem mass spectrometry (LC-MS/MS) using a Thermo Orbitrap Elite coupled with a Thermo nanoRSLC system (Thermo Fisher Scientific). Peptides were selected for fragmentation automatically by data dependent analysis. Raw data was processed using Progenesis QI (v4.1, Nonlinear Dynamics) and searched against the SwissProt and Trembl databases (accessed July 2017). Database searches were performed against the human, mouse and *E. coli* proteome databases with tryptic digest specificity, allowing a maximum of two missed cleavages, and an initial mass tolerance of 5 ppm (parts per million). Carbamidomethylation of cysteine was applied as a fixed modification, while N-acylation and oxidation of methionine, as well as the biotinylation of lysine, were applied as variable modifications during the database searches.

### Label free protein quantification and data analysis

Relative label free protein quantification of mass spectrometry experiments was performed using Progenesis QI (v4.1, Nonlinear Dynamics). Raw data was processed and aligned in Progenesis QI. Once aligned, relative protein quantification was calculated from the abundances of ions with three or more isoforms identified as being from unique (non-conflicting) peptides. Comparisons between relative protein abundances were made between proteins isolated from wild type or mutant Bid-BirA* lysates (sample) and proteins isolated from venus-BirA* lysates (control) to calculate a fold change enrichment. Sample and control protein abundances were paired based on drug treatment, e.g. WTBid-BirA* + nocodazole was compared with venus-BirA* + nocodazole. Statistical analysis of proteomics data was performed in Progenesis QI using ANOVA.

### STRING analysis

Predicted interaction networks were computed by STRING (Search Tool for the Retrieval of Interacting Genes/Proteins) from the top 10 highest scoring predicted functional partners for Bid (31, 32). Only data from experimental validated interactions and interactions in curated databases was used to score predicted functional partners. STRING Database v11.0, accessed April 2019.

### VDAC2 deletion by CRISPR/Cas9

Deletion of endogenous *VDAC2* was achieved through CRISPR/Cas9. Two sgRNA sequences (CAATGTGTATTCCTCCATC, GATGGAGGAATACACATTG) designed to target upstream of the second protein coding exon were cloned into PX458 (pSpCas9(BB)-2A-GFP). PX458 containing the VDAC2 targeting guides was transfected into low passage WT MCF-7 cells and sorted by FACS for GFP expression. Single cells were isolated from the sort and plated into individual wells of 96 well plates. Wells in which clonal cell populations had expanded were transferred to larger cultures and genomic DNA extracted for identification of indels by PCR and sequencing.

### Statistical analysis

Details of statistical tests are provided in figure legends. Statistical analysis was performed using GraphPad Prism v7.

## Acknowledgements

RP was generously supported by a studentship funded by John and Janet Hartley. LK was funded through the Manchester Cancer Research Centre by a CRUK training award. JS was funded by a Biotechnology and Biological Sciences Research Council (BBSRC, UK) David Phillips Fellowship (BB/L024551/1). VM was partially supported by a grant from the Sir Richard Stapley Educational Trust. The Wellcome Centre for Cell-Matrix Research is supported by a core grant from the Wellcome Trust (203128/Z/16/Z). We are grateful for the assistance provided by staff in the Bioimaging, Bio-MS core facilities as well as Craig Lawless for discussions relating to bioinformatics and data analysis. We would also like to thank Michael White for discussions and comments relating to the preparation of the manuscript.

## Author contributions

RP and APG conceived and planned the project. RP conducted the majority of the experiments presented in the paper. LK conducted apoptosis assays. VM and JS assisted with the analysis of the MS data. PW generated the shBid virus constructs. LK, KB and JS carried out data analysis. RP and APG wrote the paper and all authors discussed the results and commented on the manuscript.

