## Supplemental Figures 1-4 and legends for "BioID based proteomic analysis of the Bid interactome identifies novel proteins involved in cell cycle dependent apoptotic priming"

**Supplementary Figure legends.**

**Figure. S1 (Linked to Figure 1).** *Validation of a Bid-BirA* bait fusion protein.*

A. HEK-293T cells were transiently transfected with plasmids expressing either tBid-eYFP, tBid-BirA* or Venus-BirA*. After an overnight incubation cells were harvested, cytospun onto slides and stained for their respective fusion proteins tag, eYFP, venus or myc (present on the Bid-BirA*). Apoptosis was quantified by nuclear fragmentation. Data represents mean and SD of 3 independent experiments. Data analysed by one-way ANOVA, followed by Tukey's multiple comparison test. **** represents $p < 0.0001$. Scale bar = 10µm.

B. HEK293T cells transiently expressing tBid-BirA* or BirA* alone were incubated overnight in the presence or absence of 50 µM biotin. In the presence of biotin tBid-BirA* and BirA* are seen to self-label at the predicted molecular weights (indicated by red arrows). Endogenously biotinylated proteins are detected at around 280, 130, 76 and 74 kDa.

C. HEK-293T cells were transfected with plasmids expressing either GFP-Bcl-XL, or GFP-Bcl-XL and tBid-BirA*, then incubated overnight in the presence of 50 µM biotin. Biotin labelled proteins were isolated from whole cell lysates (WCL) on streptavidin beads. The WCL, the unbound fraction and the streptavidin bead bound fraction were separated by SDS-PAGE and immunoblotted for either GFP or biotin.

D. HEK-293T cells transfected and treated as in C with overnight culture in the presence or absence of 50 µM biotin as indicated. Cells were fixed and immunostained for the myc-tag and biotin. Scale bar = 10µm.

**Figure S2 (Linked to Figure 4).** *Human breast cell lines display heterogeneous responses to Taxol induced prolonged mitosis.*

A, B, C. Examples of time-lapse sequences representing: (A) Normal division; (B) Apoptosis in mitosis; (C) Mitotic slippage.

D. Single cell fate profiles of human breast carcinoma lines, MCF7 and MDA-MB-231, either control (without drug) or with 1μM Taxol, imaged over a 60 hour period. Each individual horizontal line represents a single cell. Data represents 60 cells tracked over two independent repeats.

**Figure S3 (Linked to Figure 4).** *Phosphorylation of Bid regulates apoptotic priming in mitotic breast cancer cells.*

A. Single cell fate profiles of the MCF-7 lines, untreated or treated with 1μM Taxol, in the presence of 5μM ABT-737 over 50 hours. Data represent 90 cells tracked over 3 independent experiments.

B. Summary of apoptosis in mitosis for indicated cell lines, showing that ABT737 restores the priming of shBid MCF7 cells in mitosis to that of WT cells. Data shown are mean and SD. Data were analysed by one-way ANOVA, followed by Tukey's multiple comparison test (ns = non-significant; *** = $p<0.001$).

C. Summary of duration of mitosis for indicated cell lines in B. Mean and SD plotted. Data were analysed by one-way ANOVA, followed by Tukey's multiple comparison test (ns = non-significant; **** = $p<0.0001$).

D. MDA-MB-231 cells stably expressing shBid & mouse BidWT-GFP or variant (S66A or G94E) were enriched for mitosis with overnight 18 hour treatment in nocodazole followed by shake off.

E. Single cell fate profiles of modified MDA-MB-231 lines treated with 1μM taxol over a 50 hour period. Data represents 90 cells tracked over 3 independent repeats.

**Figure S4 (Linked to Figure 5).** *VDAC2 coordinates Bid phosphorylation dependent apoptotic priming in mitosis.*

Single cell fate analysis of Wt and D11 VDAC2 KO MCF-7 cell lines. D11 cells included that are stably expressing VDAC2 V5, shBid alone or shBid in conjunction with the indicated mouse Bid-GFP variant (mBidWT-GFP, mBidS66A-GFP, mBidG94E-GFP). Cells were untreated or treated with 1μM taxol and/or 5 μM ABT-737 over 48 hours. Data represents 90 cells tracked over 3 independent experiments.

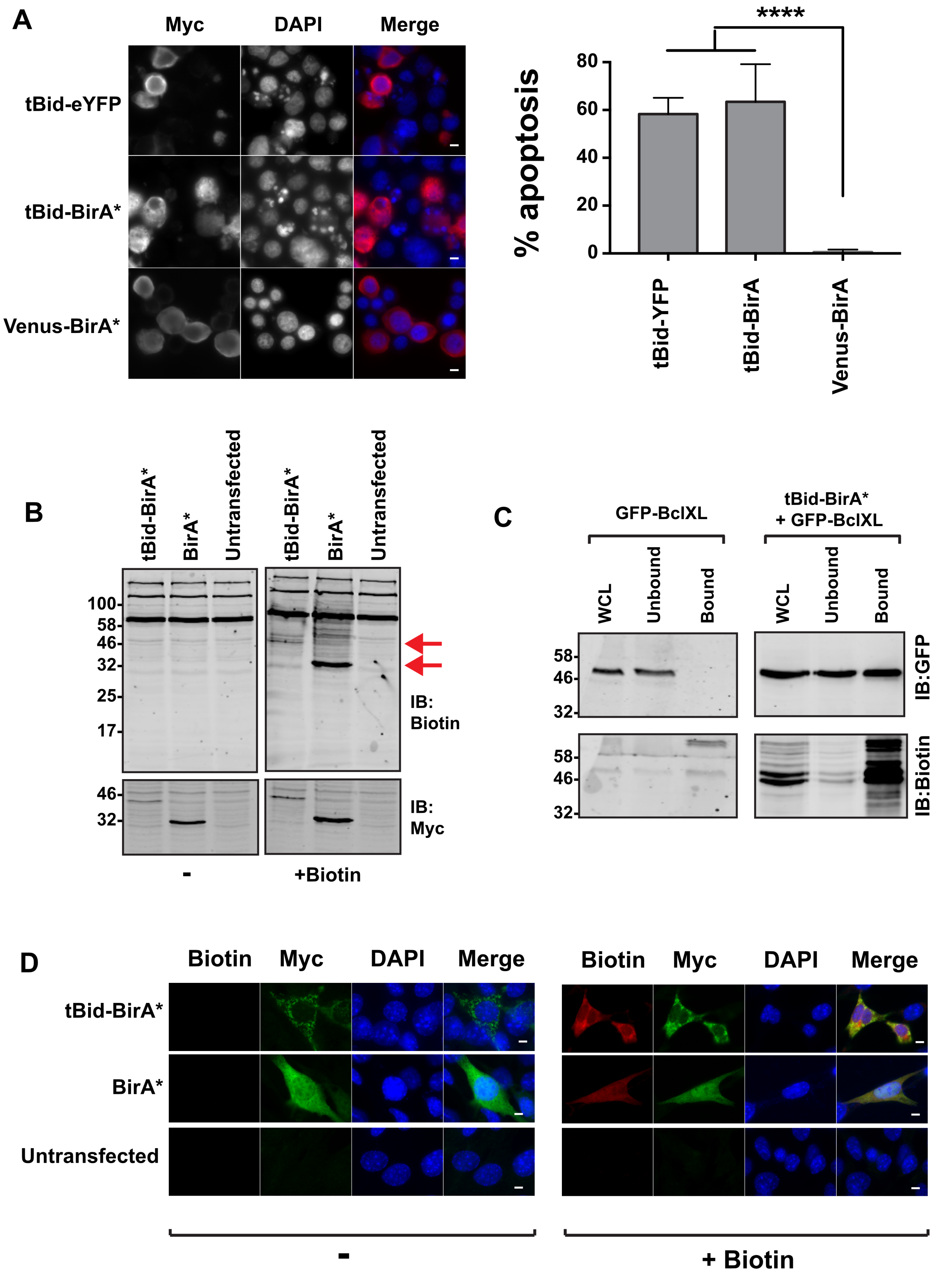


Pedley Figure S1

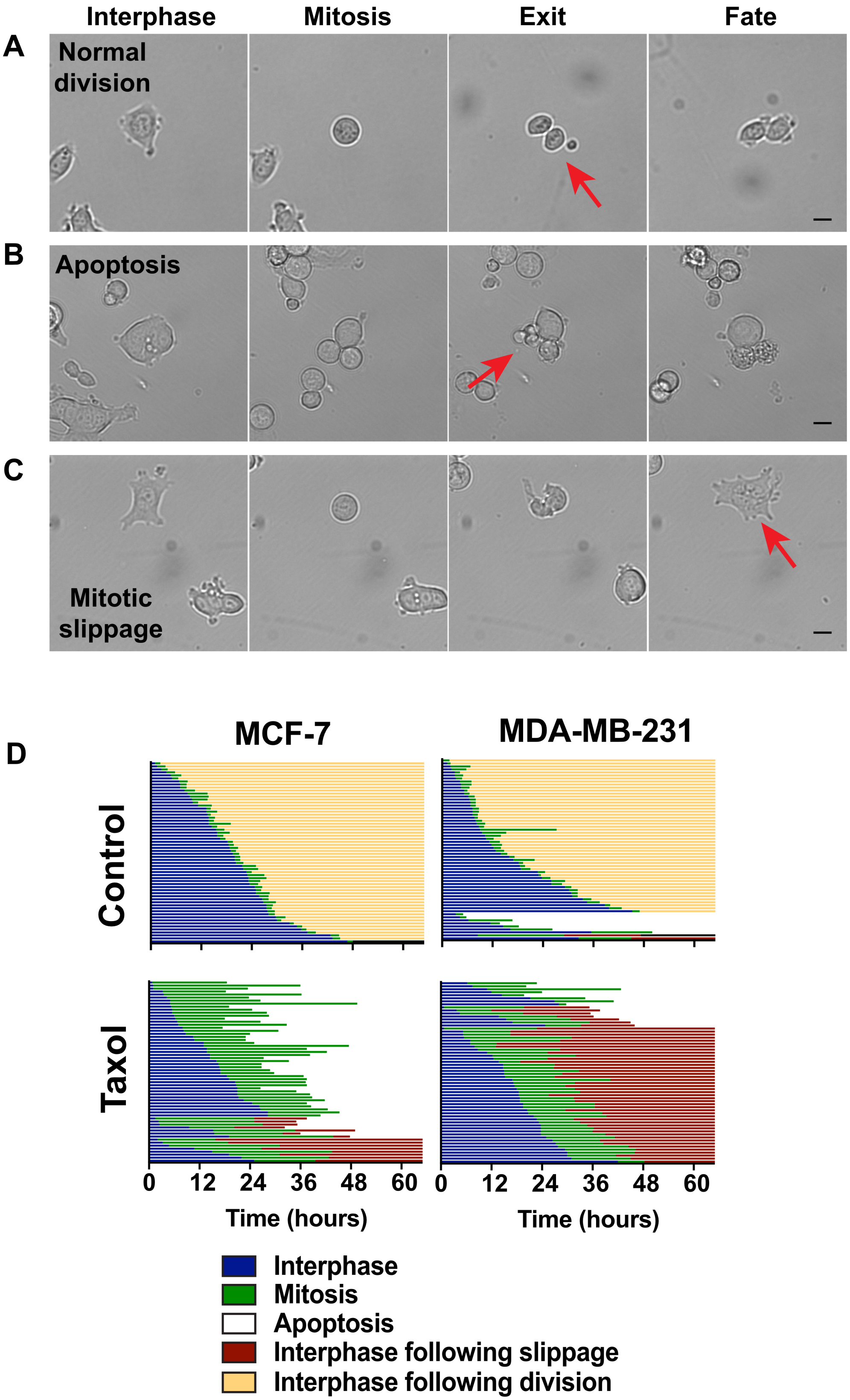


Pedley Figure S2

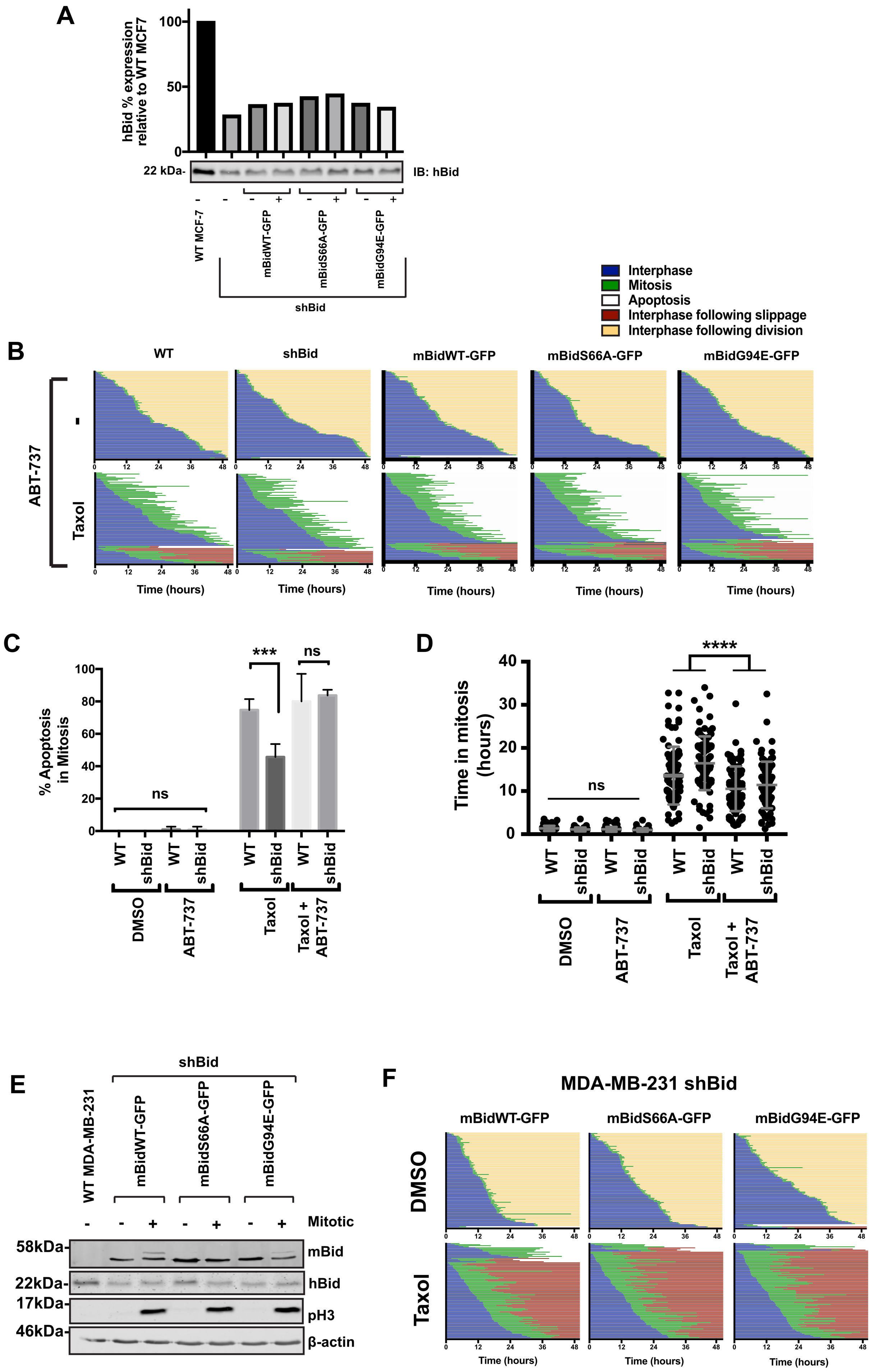


Pedley Figure S3

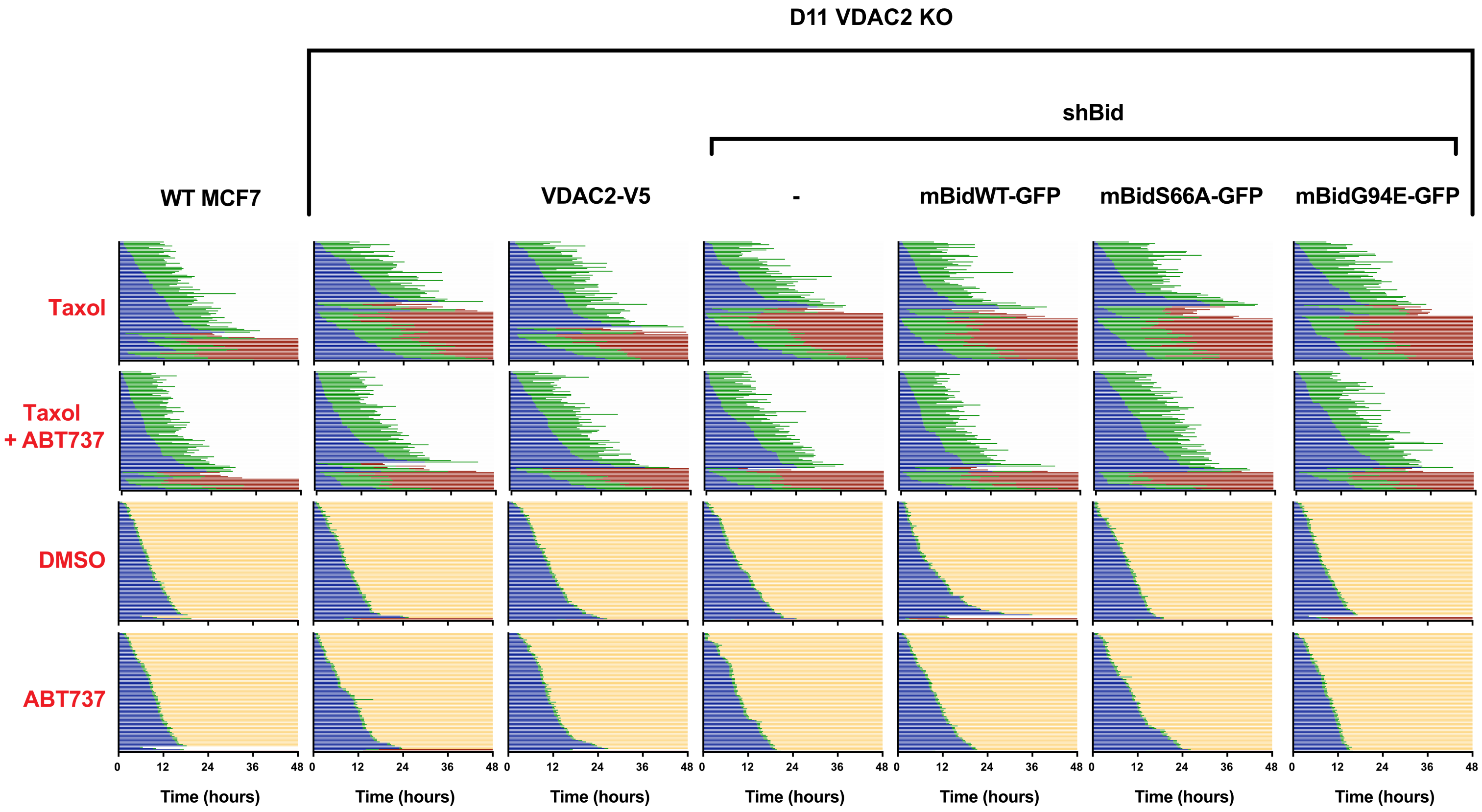


Pedley Figure S4
